# Acyl-protein thioesterase 1 (*LYPLA1*) activity promotes the growth of MDA-MB-468 triple-negative breast cancer cells

**DOI:** 10.1101/2025.10.22.683516

**Authors:** Michael Salsaa, Mahtab Tavasoli, Haggag S Zein, Shubhashree Pani, Rahul S Kathayat, Bryan C Dickinson, Gregory D. Fairn

**Affiliations:** Department of Pathology, Dalhousie University, Halifax, Nova Scotia, Canada; Department of Biochemistry & Molecular Biology, Dalhousie University, Halifax, Nova Scotia, Canada; Department of Pharmacology, Dalhousie University, Halifax, Nova Scotia, Canada; Department of Chemistry, University of Chicago, Chicago, United States; Chan Zuckerberg Biohub, Chicago, IL 60642

## Abstract

Protein *S-*acylation is a lipid-based, often reversible post-translational modification that can regulate many aspects of protein behavior, including subcellular localization, protein-interactions, and activity. Emerging evidence has identified roles for individual protein acyltransferases encoded by the *ZDHHC* in cancers, yet the roles of de-*S-*acylation enzymes are less clear. Recent evidence suggests that acyl-protein thioesterase (APT1)/*LYPLA1* can impact epithelial-mesenchymal transition and metastasis. This study integrates patient datasets, CRISPR dependency data, and *in vitro* assays to find APT1 as a context-dependent vulnerability in triple-negative breast cancer (TNBC). Despite the highest protein abundance in luminal MCF7 cells, basal-like MDA-MB-468 cells exhibited the most prominent specific APT1 activity, reflecting subtype-specific regulation. Inhibition of APT1 with ML348 increased *S*-acylation of nuclear and mitochondrial proteins without altering global acylation. Functionally, APT1 inhibition reduced cell proliferation while inducing minimal apoptosis, consistent with cytostatic growth arrest. Cell-cycle analysis revealed G1 accumulation and reduced S/G2 transition, linking proteomic changes to impaired replication. These findings establish APT1 as a regulator of TNBC proliferation through dynamic de-*S-*acylation of cell-cycle and mitochondrial proteins, highlighting it as a potential therapeutic vulnerability in aggressive breast cancers.

## Introduction

Breast cancer is one of the most commonly diagnosed cancers and the leading cause of cancer-related death among women worldwide ^1,2^. It is a heterogeneous disease comprising distinct molecular subtypes, with each characterized by unique biological features and clinical outcomes ^3,4^. Among these, triple-negative breast cancer (TNBC) is defined by the absence of estrogen receptor (ER), progesterone receptor (PR), and HER2 expression ^5^. The lack of these common therapeutic targets limits treatment options for TNBC. Currently, the median survival for patients with TNBC remains poor, typically not exceeding five years ^6,7^. These challenges highlight the need to identify vulnerabilities and develop targeted therapies for this aggressive subtype.

Post-translational modifications (PTMs) are critical in regulating key metabolic and cellular functions by influencing protein localization, stability, folding, and enzymatic activity ^8^. Large-scale analyses across cancer types have identified common PTM signatures associated with cancer hallmarks, underscoring their therapeutic potential ^9–11^. One such modification, protein *S-*acylation, involves the covalent attachment of a fatty acid—commonly palmitate—to cysteine residues, thereby increasing the hydrophobicity of the protein ^12^. This modification impacts target proteins in many ways, such as promoting membrane association, sorting and subcellular trafficking, protein stability, and activity ^13^. The dynamic nature of *S-*acylation is controlled by opposing enzymatic activities: zDHHC acyltransferases that add fatty acids to proteins, and thioesterases that remove them ^13^. Acyl-protein thioesterase 1 (APT1), encoded by the *LYPLA1* gene, is well established as an enzyme responsible for de-*S-*acylation, or simple deacylation, of target proteins ^14–18^.

A growing body of evidence implicates APT1 in cancer progression through its regulation of protein deacylation, affecting several key pathways across tumor types. APT1 promotes epithelial–mesenchymal transition (EMT), invasion, and migration—hallmarks of metastasis ^19–23^. In cervical cancer cells (HeLa, CaSki, and SiHa), APT1 deacylation of Flotillin-1 prevents the internalization and lysosomal degradation of the insulin-like growth factor-1 receptor (IGF-1R). The resulting increase in IGF-1 signaling promotes EMT and invasive behavior ^19^. In non-small cell lung cancer, APT1 facilitates EMT via an undefined mechanism ^20^. APT1 also enhances invasiveness in melanoma, where it deacylates MUC18/melanoma-cell adhesion molecule (MCAM), increasing cell invasion ^21^. In addition to enhancing motility and invasiveness, APT1 also contributes to tumor cell survival. In chronic lymphocytic leukemia, APT1 deacylates the death receptor CD95/Fas, impairing its ability to initiate apoptosis in response to Fas ligand (CD95L) ^22^. These studies suggest that APT1 contributes to malignancy through diverse protein substrates and mechanisms that converge on enhanced survival, migration, and EMT, highlighting its potential as a therapeutic target in multiple cancer contexts.

Despite growing evidence linking APT1 to cancer progression, its gene expression, enzymatic activity, and functional relevance across breast cancer subtypes—particularly TNBC—remain poorly characterized. The potential role of APT1 in supporting the proliferation of aggressive TNBC cells has not been thoroughly explored. In this study, we examined the prognostic significance and functional dependency of APT1 across breast cancer subtypes using publicly available datasets and *in vitro* models. We analyzed APT1 protein expression and enzymatic activity in representative breast cancer cell lines and assessed the impact of its pharmacological inhibition using the selective inhibitor ML348 ^24^. Our findings reveal subtype-specific differences in APT1 activity and identify a previously unrecognized role for APT1 in sustaining TNBC proliferation, in part by modulating cell cycle progression.

## Materials and methods

### Cell Culture and Treatment

Human breast cancer cell lines MDA-MB-468 (ATCC HTB-132) and MCF7 (ATCC HTB-22) were obtained from ATCC. MDA-MB-231 and SKBR3 cells were generously provided by Dr. Paola Marcato (Dalhousie University). Cells were maintained in Dulbecco’s Modified Eagle Medium (DMEM, Wisent) supplemented with 5% fetal bovine serum (Wisent) and cultured in a humidified incubator at 37°C with 5% CO□. Once cultures reached approximately 70% confluency, cells were treated with either DMSO (vehicle control) or 50 µM ML348 (Cayman Chemical) for the indicated time periods. Cells were routinely tested for mycoplasma using a PCR-based assay.

### APT1 activity

For cellular/mitochondrial APT1 activity, cells were incubated with either 10□µM DPP-3 or 2.5□µM mitoDPP-3 in Hanks’ Balanced Salt Solution (HBSS, Wisent) for 30 minutes ^25^. Fluorescence was then measured using flow cytometry (BD LSR Fortessa™; 488 nm laser with 530/30 nm emission filter). To evaluate APT1 activity in cell lysates, cells were harvested and lysed in a non-denaturing buffer (1% NP-40, 150□mM NaCl, 1□mM EDTA, 50□mM HEPES [pH 7.4], 1□mM PMSF, and 1X cOmplete protease inhibitor cocktail [Roche, 11836145001]). Nuclei were removed by centrifugation at 17,000 x *g* for 15 minutes at 4° C. The resulting lysates were serially diluted in APT1 activity buffer (20□mM HEPES [pH 7.4], 150□mM NaCl, 0.1% Triton X-100), and protein concentrations were determined using a BCA assay (ThermoFisher). APT1 activity was assessed using a SpectraMax® i3x in a 96-well plate across multiple protein concentrations, each measured in technical duplicates, by adding 10□µM DPP-3 and recording fluorescence over 20 minutes. Specific activity was calculated from the slope corresponding to the protein concentration range that maintained linearity of the reaction.

### Western blot analysis

Cells were cultured as described above and subsequently harvested. Cell pellets were lysed in RIPA buffer without SDS, and nuclei were removed by centrifugation at 17,000 x*g* for 10 minutes. The supernatants were mixed with 4X SDS loading buffer, separated on a 10% SDS-PAGE gel, and then transferred to a polyvinylidene difluoride (PVDF) membrane. The membrane was incubated with a primary antibody against Apt1/Lypla1 (Proteintech, 16055-1-AP; 1:2000), followed by an IR700-conjugated fluorescent secondary antibody (AzureSpectra, AC2134; 1:10000). Signal was detected (Ex/Em 690/709 nm) using an Azure Biosystems 600 imaging system, and band intensities were quantified with AzureSpot Pro software. No-Stain™ protein labeling reagent (Invitrogen, A44717, Ex/Em 488/590 nm) was used to normalize for protein loading.

### Cell cycle analysis

Cells were treated as described above, trypsinized, and counted. Cell concentration was adjusted to 1 × 10□ cells/mL. Cells were pelleted and resuspended in 50 µL of HBSS containing 2% FBS (Wisent). Fixation was achieved by slowly adding 1 mL of ice-cold 70% ethanol with gentle vortexing, followed by incubation at 4°C overnight. After fixation, cells were pelleted and washed twice with PBS, then stained with 0.1 mg/mL DAPI in PBS containing 0.1% Triton X-100. Following a 30-minute incubation at room temperature, fluorescence was measured using a BD LSRFortessa™ flow cytometer (Ex/Em 405/460 nm), and cell cycle analysis was performed using the built-in functionality of FlowJo™ software.

### Quantifying Live/Dead Cells

Live cells were quantified by staining with 250 nM Calcein AM (Cayman Chemical) for 15 minutes. Following staining, cells were washed with PBS, trypsinized, and collected. The cell pellet was then resuspended in 100 µL of Annexin-V Binding Buffer (BioLegend®, 422201) with 3 µL of Annexin-V-APC (BioLegend®, 640920) to label apoptotic cells. To differentiate apoptotic from necrotic cells, 0.5 µL of 1000x 7-AAD (Cayman Chemical, 400201) was added, and the cells were incubated for an additional 15 minutes. Fluorescence was subsequently measured using a BD LSRFortessa™ flow cytometer, and data were analyzed with FlowJo™ Software.

### Cell Proliferation

MDA-MB-468 cells were seeded at 5 × 10□ cells per well in a 96-well plate and incubated overnight. One day post-seeding, cell growth was measured in a subset of wells using a colorimetric WST-8 assay (Cayman Chemical, #10010199) following the manufacturer’s instructions. The remaining wells were then treated with DMSO, 50 µM ML348, or switched to starvation media for 24 or 48 hours, and cell proliferation was assessed as described above.

### Acyl-Resin Assisted Capture (RAC)

MDA-MB-468 cells were treated with 50 µM ML348 for 24 hours, harvested, and lysed in a buffer containing 1% NP-40, 150 mM NaCl, 1 mM EDTA, 50 mM HEPES (pH 7.4), 1 mM PMSF, and 1X cOmplete protease inhibitor cocktail (Roche, #11836145001). Nuclei were removed by centrifugation. The lysates were processed as described ^26^. Briefly, each 100 µL of lysate was incubated with 300 µL of blocking buffer (2.5% SDS, 100 mM HEPES, 1 mM EDTA) and 4 µL of *S-*methyl thiomethanesulfonate (MMTS; Sigma-Aldrich, #208795) for 1 hour at 40°C with constant shaking at 1000 rpm. Proteins were precipitated with 1.1 mL 100% acetone, and the pellets were washed with 70% acetone to remove excess MMTS. The pellets were then resuspended in binding buffer (100 mM HEPES [pH 7.4], 1 mM EDTA, 1% SDS) by sonication. Hydroxylamine (NH□OH, from Sigma-Aldrich, 0.6 M final concentration) or NaCl (for -HAM control samples) was added, followed by an end-over-end rotator incubation with 30 µL of packed acyl-RAC beads (High-Capacity Acyl-RAC S3 beads; Nanocs, #AR-S3-1,2) for 2 hours at room temperature. Beads were subsequently washed three times with binding buffer and twice with 50 mM Tris-HCl (pH 7.5).

For western blot analysis, SDS loading buffer containing 10% β-mercaptoethanol was added, and samples were briefly boiled before being loaded onto a gel. After transfer, membranes were stained with a No-stain protein labeling reagent (Invitrogen, A44717) following the manufacturer’s instructions. Alternatively, for mass spectrometry analysis, on-bead digestion was performed using trypsin/Lys-C (1 µg per sample in 50 mM NH□HCO□, pH 8.3) for 5 hours at 37°C on an end-over-end rotator. Beads were rinsed with 50 mM NH□HCO□ (pH 8.3), and the supernatant fractions were pooled, desalted, and lyophilized prior to LC-MS analysis.

### Mass Spectrometry

Peptide samples were analyzed using an Orbitrap Lumos mass spectrometer (Thermo Fisher Scientific) coupled to an M-Class UPLC system (Waters). Peptides were loaded onto a C18 trap column (180 µm × 20 mm, 5 µm) and separated on an analytical C18 column (75 µm × 250 mm,1.7 µm) with a 60 min gradient from 1% to 30% solvent B (0.1% formic acid in acetonitrile) at a flow rate of 300 nL/min. Solvent A was 0.1% formic acid in water.

Data were acquired in data-dependent acquisition mode with full MS scans (m/z 400–1600) at 120,000 resolution in the Orbitrap, followed by HCD MS/MS of the most intense precursors (charge 2–4) at 30% normalized collision energy and 30,000 resolution. The isolation window was set to 1.6 m/z, and dynamic exclusion was applied for 60 s.

Raw data were processed using FragPipe (v23.0) with MSFragger (v4.3) ^27^ against the *Homo sapiens* UniProt database. Searches were performed with precursor and fragment tolerances of 20 ppm, carbamidomethylation of cysteine as a fixed modification, and oxidation of methionine and N-terminal acetylation as variable modifications. Peptide spectral matches were rescored with MSBooster and validated with Percolator, and protein inference was performed using ProteinProphet with a false discovery rate (FDR) < 1% at the PSM, peptide, and protein levels. Label-free quantification was carried out with IonQuant using default settings. Processed data were analyzed in R for statistical and visualization purposes.

## Results

### LYPLA1 in breast cancer: prognostic significance and cellular fitness

To evaluate the prognostic usefulness of *LYPLA1* in breast cancer, we performed a Kaplan-Meier survival analysis on a cohort of 4,929 patients, dichotomized by median gene expression ^28^. As shown in Figure 1A, patients with low *LYPLA1* expression (black line) exhibited a significantly higher overall survival than those with high expression (red line). The median survival was 216.66 months for the low expression group versus 191.21 months for the high expression cohort. The hazard ratio of 1.18 (95% CI: 1.07–1.31, p = 0.0013) indicates that high *LYPLA1* expression is associated with an 18% increased risk of death, supporting its role as a potential prognostic biomarker.

**FIGURE 1.**
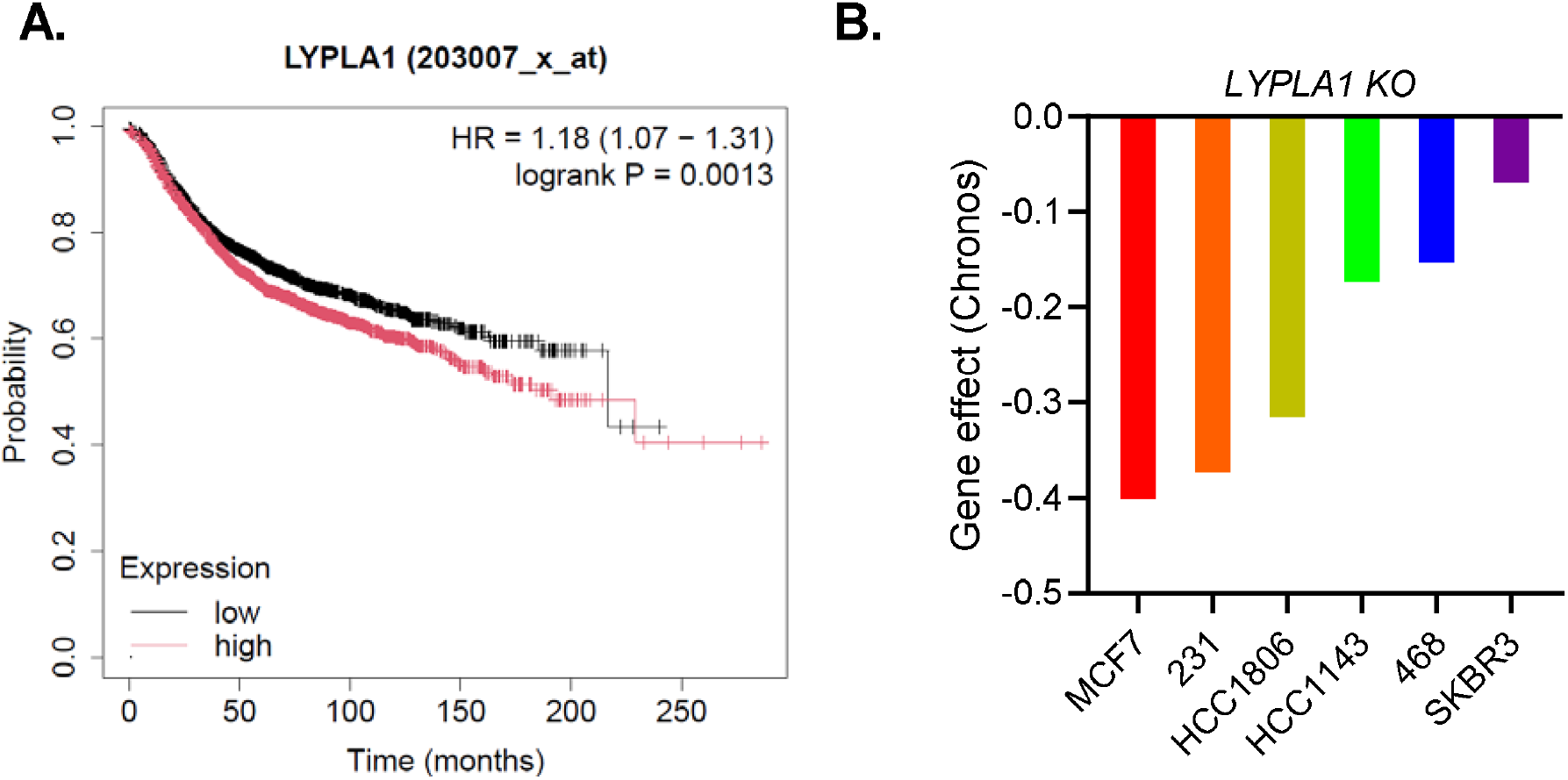
Impact of *LYPLA1* on Breast Cancer phenotypes. A, Kaplan-Meier survival analysis of breast cancer patients stratified by *LYPLA1* expression. Censoring is marked by tick marks along the curves. B, Bar graph displaying the *LYPLA1* CRISPR dependency scores (Chronos scores) for the indicated breast cancer cell lines, derived from the DepMap Public dataset. More negative Chronos scores indicate a greater dependency on *LYPLA1* for cell viability, suggesting that its knockout impairs cell growth or viability. Each bar represents a distinct breast cancer cell line, highlighting the variability of *LYPLA1* dependency across different subtypes.

The Cancer Dependency Map (DepMap) is a comprehensive resource that elucidates the genetic dependencies of cancer cells using large-scale datasets, such as CRISPR-Cas9 knockout screens, to identify genes essential for cell survival and proliferation (https://depmap.org/portal). To assess the impact of *LYPLA1* deletion on breast cancer cells, we queried the DepMap public database ^29^. Figure 1B displays the Chronos scores for several breast cell lines following CRISPR-mediated knockout of *LYPLA1*. The Chronos score quantifies gene essentiality, where more negative values indicate a greater dependency on the gene for cell viability. In this analysis MDA-MB-231 (231), HCC1806, HCC1143, and MDA-MB-468 (468) are TNBC cell lines.

### APT1 protein levels and activity across Breast Cancer subtypes

To investigate the variability in Chronos dependency scores for *LYPLA1*, we assessed its protein product, APT1, across representative breast cancer subtypes using Western blot analysis (Figure 2A). We selected cell lines that model distinct BC subtypes: MCF7, a luminal A cell line (ER^+^/PR^+^/HER2^−^); SKBR3, a HER2-enriched line (ER^−^/PR^−^/HER2^+^); and MDA-MB-231 (231) and MDA-MB-468 (468), which are triple-negative breast cancer (TNBC) cell lines (ER^−^/PR^−^/HER2^−^). Among these, MCF7 exhibited the highest APT1 protein levels, followed by 468. In contrast, SKBR3 and 231 displayed comparably lower APT1 expression (Figure 2B).

**FIGURE 2.**
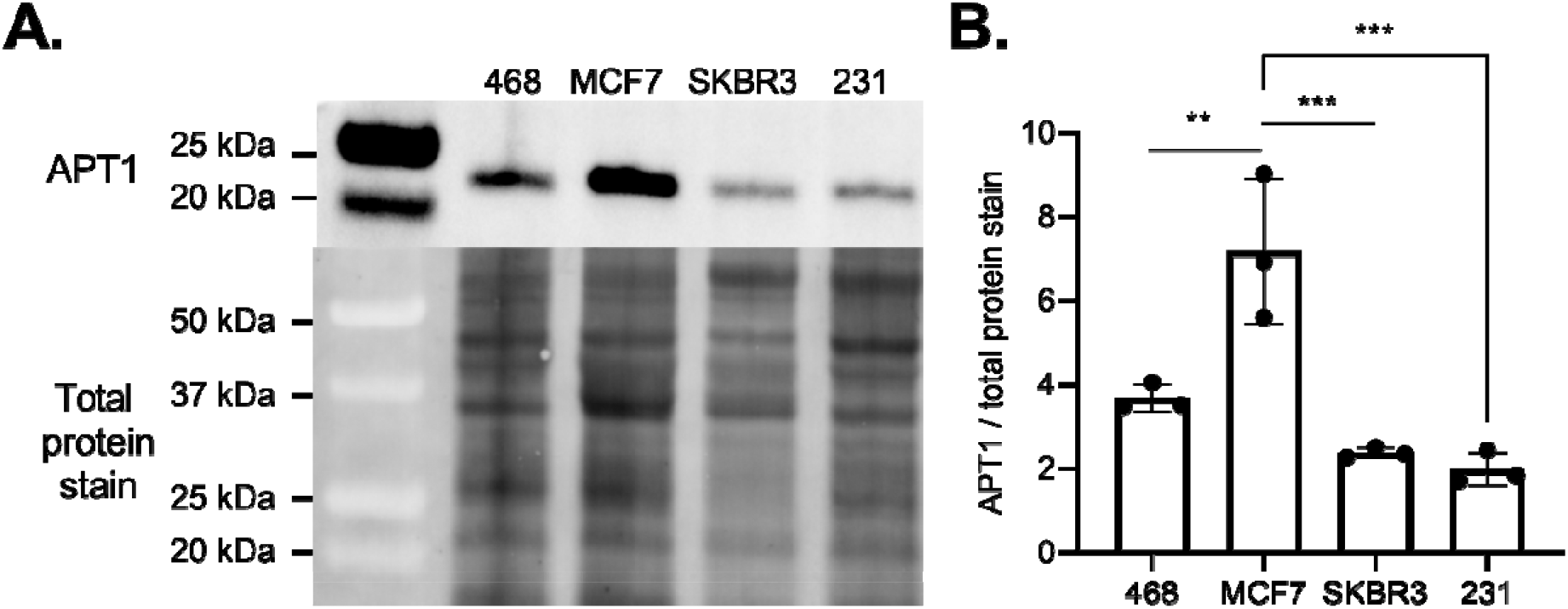
Breast cancer cell lines have varying levels of APT1. A, representative Western blot analysis of APT1 protein levels in several breast cancer cell lines. Total protein staining (No-Stain™ protein labeling reagent) was used to normalize for protein loading. B, quantification of APT1 levels using AzureSpot Pro image analysis software. Data shown are mean ± SD (n=3). Statistical significance was determined using one-way ANOVA followed by Tukey’s multiple comparisons test (** p < 0.01, *** p < 0.001).

APT1 was originally characterized as a cytosolic enzyme with multiple protein substrates ^30^. However, more recent studies have shown that APT1 is predominantly localized to mitochondria ^25,31^, although its mitochondrial function remains poorly understood ^32^. To explore whether BC subtypes differ in cytosolic and mitochondrial APT1 activity, we quantified APT1 activity in both compartments.

To assess cytosolic APT1 activity, we used the fluorogenic probe DPP-3, which remains non-fluorescent until selectively cleaved by APT1 ^33^. Upon cleavage, the probe emits a fluorescent signal that can be quantified via flow cytometry. Cytosolic APT1 activity mirrored protein expression levels across the cell lines, with MCF7 displaying the highest activity, followed by 468. In contrast, SKBR3 and 231 exhibited lower APT1 activity (Figure 3A). Mitochondrial APT1 activity was evaluated using a mitochondria-targeted version of the probe, mitoDPP-3 ^25^. Fluorescence in 468 cells was >4-fold higher than in the other cell lines (Figure 3B), indicating substantially elevated mitochondrial APT1 activity.

**FIGURE 3.**
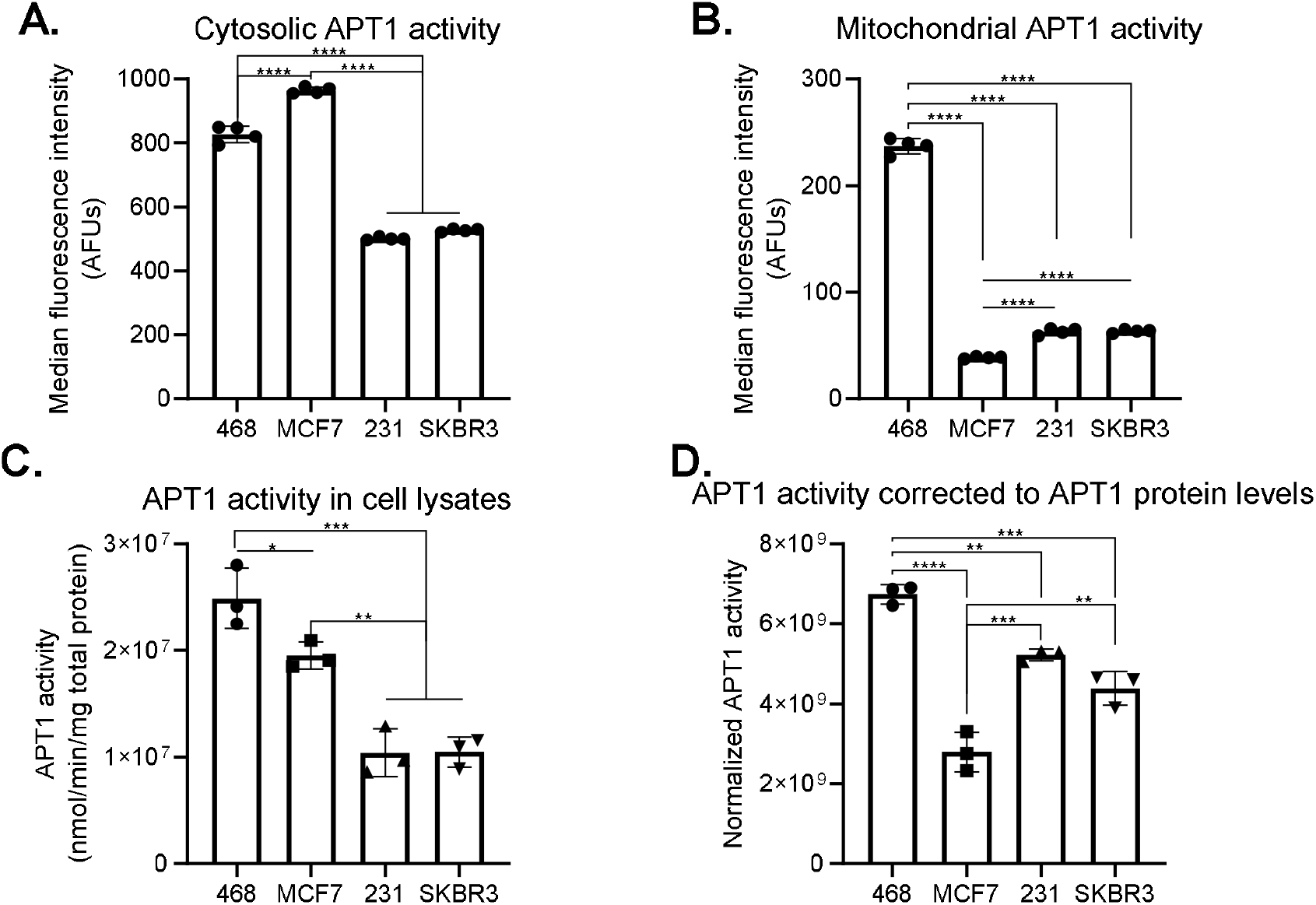
MDA-MB-468 displays the highest APT1 activity. A, cytosolic and B, mitochondrial APT1 activity in cancer cell lines were assayed using DPP-3 and mito-DPP3 fluorescence, respectively. Fluorescence was measured using flow cytometry. Median fluorescence intensity (MFI) of single cells was analyzed with FlowJo. Data shown are mean ± SD (n=4). C, APT1-specific activity was assayed in cell lysates. DPP3 fluorescence (Δ λ_em_ 545/20□nm/min/mg protein) was measured using a SpectraMAX i3X plate reader at three protein concentrations to ensure linearity of the reaction. Data shown are mean ± SD (n = 3, with at least two technical replicates). D, APT1-specific activity was normalized to APT1 levels as detected by WB analysis in Fig. 2. Statistical significance was determined using one-way ANOVA followed by Tukey’s multiple comparisons test (** p < 0.01, *** p < 0.001, **** p < 0.0001).

Direct comparison of the signals from DPP-3 (cytosol) and mitoDPP-3 (mitochondria) is not possible because the two probes measure activity in distinct subcellular compartments with different volumes, geometries, and uptake mechanisms. In addition, mitoDPP-3 uptake depends on mitochondrial membrane potential, introducing a variable with no counterpart in the cytosolic assay. To eliminate these confounding factors, we lysed the cells under non-denaturing conditions—thereby collapsing membrane potential and mixing all compartments—and measured DPP-3 fluorescence in the resulting whole-cell lysates. In this cell-free assay, 468 cells displayed the highest APT1-dependent signal (Figure□3C), indicating elevated enzymatic activity that cannot be explained by protein abundance alone. To quantify specific APT1 activity, we normalized the lysate fluorescence to APT1 protein levels determined in Figure□2, generating an activity-per-protein index (Figure□3D). Normalization confirmed 468 as the most active cell line and also revealed that the high APT1 abundance in MCF7 translates to a comparatively low catalytic activity, whereas 231 and SKBR3 exhibit higher specific activity than MCF7.

### APT1 inhibition by ML348 results in selective S-acylation changes, not global remodeling

We selected the 468 cell line for further analysis due to its high APT1 enzymatic activity. Because APT1 is a protein thioesterase, its inhibition is expected to increase *S*-acylation of target proteins. To test this, 468 cells were treated with ML348—a reversible and selective APT1 inhibitor—for 24 hours, and *S*-acylated proteins were isolated using the acyl-resin–assisted capture (acyl-RAC) method. In this approach, free cysteine residues are first blocked with MMTS, acyl–cysteine thioester bonds are cleaved with hydroxylamine, and the resulting free thiols are captured on sulfhydryl-reactive pegylated resin through mixed disulfide exchange ^34^. This assay can be readily combined with protein stains and mass spectrometry for protein detection and quantification.

We first examined the highly abundant proteins separated by SDS-PAGE. Total protein levels were visualized using the No-Stain™ protein labeling reagent, and *S*-acylated proteins in the hydroxylamine-treated fraction were normalized to total protein levels in the input fraction. ML348 treatment did not produce a detectable increase in overall *S*-acylation of highly abundant proteins compared to control-treated cells (Figure 4), suggesting that APT1 inhibition does not induce marked alterations in the steady-state levels of highly abundant proteins in 468 cells. Therefore, we proceeded with mass spectrometry analysis to identify proteins with more subtle alterations in *S*-acylation. Mass spectrometry analysis using intensity-based Absolute Quantification (iBAQ)-^35–37^ identified a distinct set of proteins with increased *S*-acylation following ML348 treatment. The distribution of these changes is shown in a volcano plot (Figure 5). Given the primarily mitochondrial localization of APT1, we found multiple mitochondrial proteins—such as *ACOT9, CYC1, SLC25A3*, and *MTCH2*—suggesting effects on mitochondrial metabolism and bioenergetics ^38–41^. Additionally, we identified several nuclear proteins that showed enhanced *S*-acylation following APT1 inhibition, including *MCM2, TK1, DHX9, RBBP4*, and *RBBP7*, which are involved in replication licensing, nucleotide metabolism, RNA processing, and chromatin regulation, respectively ^42–46^. These findings indicate that APT1 inhibition alters nuclear and mitochondrial acylation profiles, revealing diverse cellular processes potentially influenced by APT1 activity.

**FIGURE 4.**
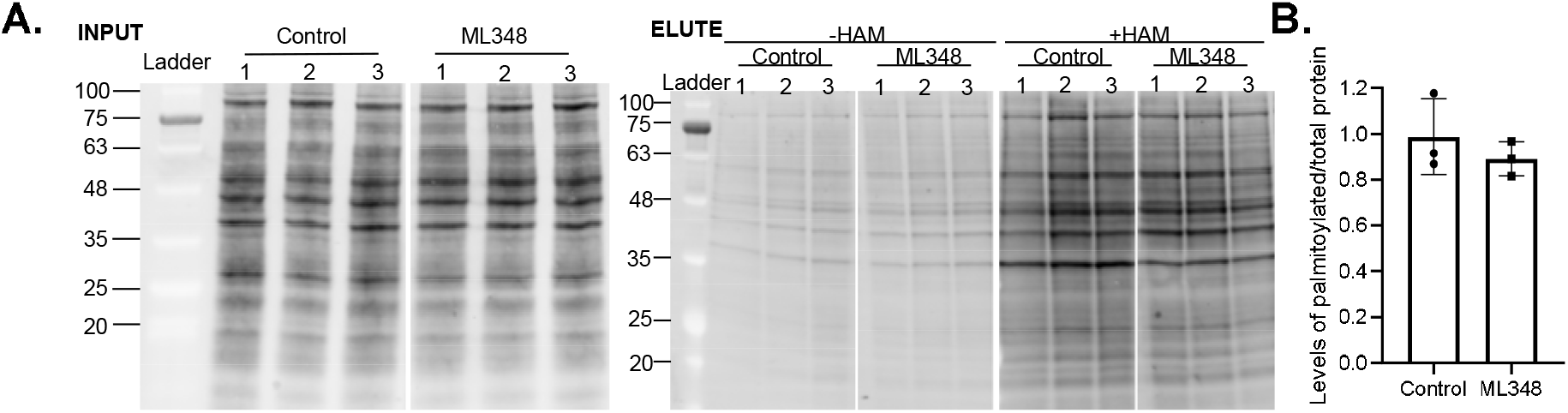
ML348 does not alter overall *S*-acylated protein levels. A, 468 cells were treated with 50 µM ML348 for 24 hours. Cell lysates were subjected to acyl-RAC followed by Western blot analysis. The input represents total cell lysate collected before acyl-specific enrichment. Half of the lysate was treated with 0.6 M hydroxylamine (+HAM) to cleave *S*-acyl modifications. In contrast, the other half was incubated with 0.6 M NaCl (−HAM) as a control for nonspecific bead binding. Total protein was visualized using the No-Stain™ reagent. B, Quantification of the results in A; total *S*-acylated protein levels in the +HAM condition were normalized to the corresponding total protein levels in the input. Data are presented as mean ± SD (n = 3).

**FIGURE 5.**
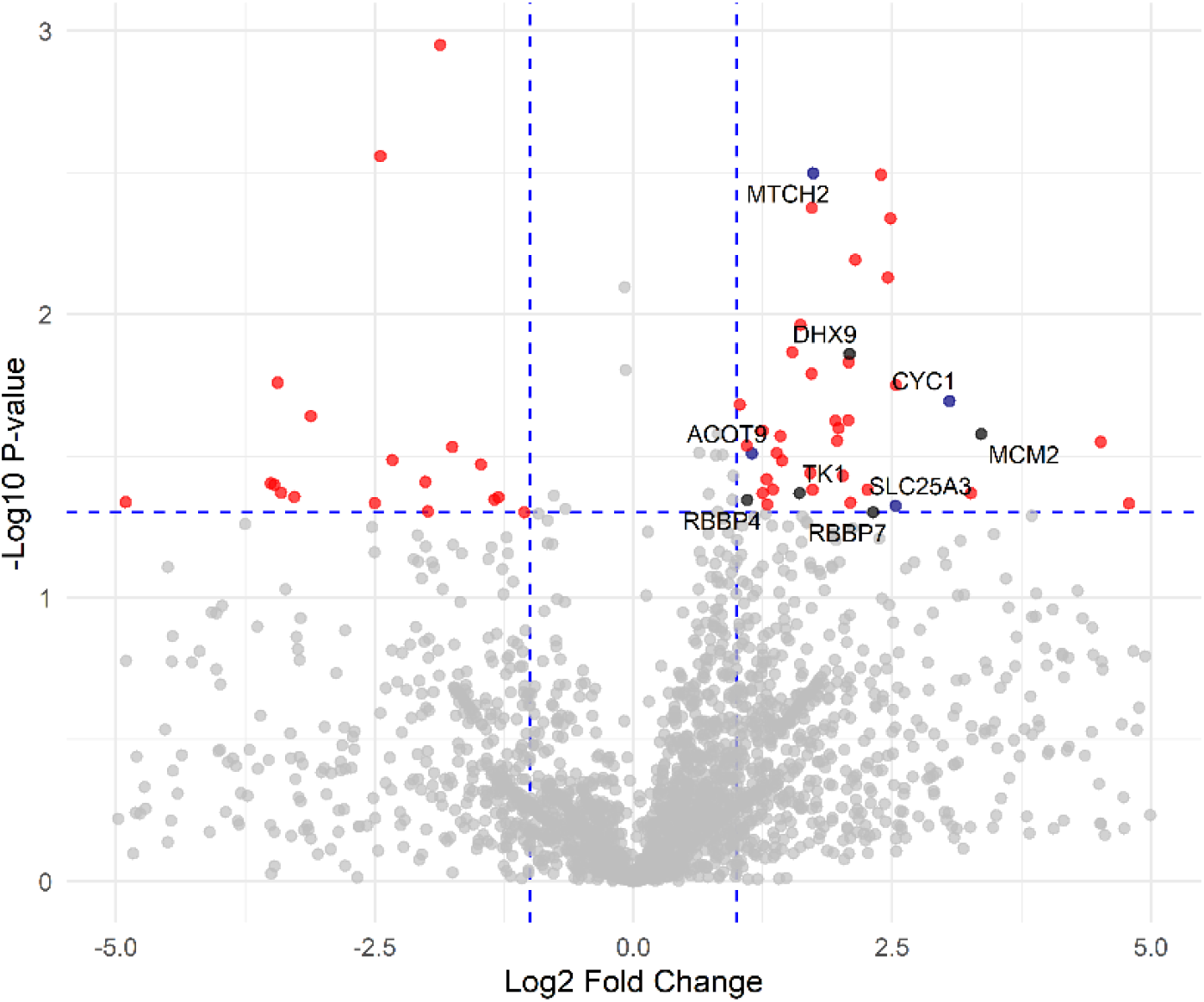
Volcano plot of proteins with altered *S*-acylation following APT1 inhibition. Volcano plot of log□ fold change versus –log□□ p-value from acyl-RAC mass spectrometry analysis (n = 5, unpaired t-test). Dashed lines indicate thresholds for significance (log□FC ≥ |1|; p < 0.05). Significant proteins are shown in red, non-significant in grey, with mitochondrial proteins highlighted in navy blue and cell cycle/transcription-associated proteins in black.

### ML348 decreases cell proliferation without inducing cell death

Given the enrichment of nuclear and mitochondrial proteins following APT1 inhibition, we next sought to determine whether these molecular changes affected cell proliferation. To this end, we used the WST-8 (also known as CCK-8) assay, which measures the reduction of the tetrazolium salt WST-8 by cellular dehydrogenases to generate a water-soluble formazan dye ^47^. The absorbance of this dye reflects overall metabolic activity and serves as a proxy for viable cells. Treatment with ML348 significantly reduced cell viability at 24 and 48 hours to levels comparable to those observed under serum-starved conditions (Figure 6A).

**FIGURE 6.**
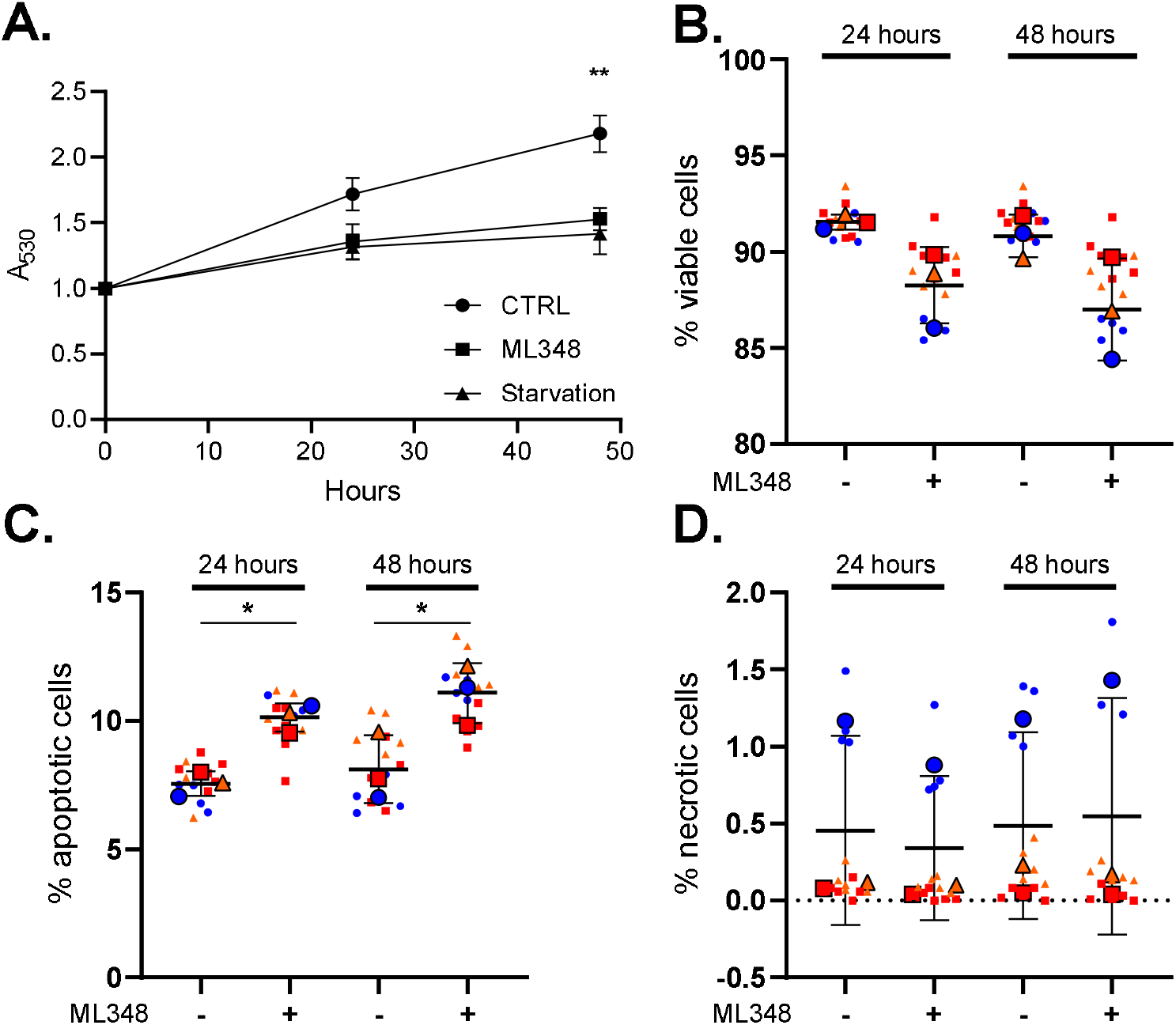
ML348 decreases MDA-MB-468 proliferation while minimally impacting cell death. A, Cell proliferation was assayed using WST-8 cell proliferation assay. Cells were plated in a 96 well-plate at a density of 10^4^ cells per well. After overnight incubation of the plate, a baseline for cell growth was established by measuring formazan absorbance at 530 nm. Cells were then either treated with vehicle, 50 µM ML348, or switched to DMEM with no FBS to establish a starvation condition. After 24 and 48 hours of treatment, cell proliferation was assayed. Results were normalized to the established baseline for each replicate and corrected to 1. Statistical significance was established using Two-way repeated-measures ANOVA with Tukey’s post-hoc test. Data shown are mean ± SD (n = 3, with at least six technical replicates, ** p < 0.01). B,C,D, Cells were treated with 50 µM ML348 for 24 or 48 hours and subsequently assessed by flow cytometry using Calcein AM, Annexin-V-APC, and 7-AAD to identify (B) live,

Since ML348 treatment significantly reduced cell abundance, we next investigated whether this was due to increased cell death. Given the enrichment of mitochondrial proteins in our acyl-proteomics dataset, we reasoned that APT1 inhibition could potentially disrupt mitochondrial integrity and trigger apoptosis. To evaluate this, cells were stained with Calcein AM (viable cells), Annexin V (apoptotic marker), and 7-AAD (necrotic marker) following 24 and 48 hours of ML348 treatment. Cells positive for both Calcein AM and Annexin V were classified as early apoptotic, whereas double-positive Annexin V/7-AAD cells were considered necrotic.

ML348 treatment did not produce a statistically significant reduction in the proportion of viable cells (Figure 6B). Although ML348 treatment significantly increased the proportion of apoptotic cells, the overall change was minimal—from 7.5% to 10% at 24 hours and from 8% to 11% at 48 hours (Figure 6C). Necrotic cell death remained negligible (<1%) in both treated and control conditions (Figure 6D). The low frequency of apoptotic populations indicates that APT1 inhibition does not cause substantial mitochondrial damage, despite the enrichment of mitochondrial proteins observed in the acyl-proteomics dataset. Consequently, the reduction in WST-8 signal primarily reflects decreased cell proliferation rather than extensive cell death, supporting its validity as a proxy for total cell number in this context. Together, these findings suggest that APT1 inhibition suppresses cell growth mainly by impairing proliferative capacity rather than by inducing cytotoxicity.

(C) apoptotic, and (D) necrotic cells, respectively. Data is presented as the percentage of cells within the gated single-cell population. Statistical significance was determined using two-way ANOVA with Sidak’s post-hoc test. Results are expressed as mean ± SD (n = 3, with at least four technical replicates, * p < 0.05).

### ML348 induces G1 arrest and impairs cell cycle progression

Proteomic analysis revealed increased *S*-acylation of nuclear proteins involved in replication and chromatin regulation, suggesting that APT1 inhibition could perturb pathways required for DNA synthesis and cell-cycle control. Consistent with this, ML348 treatment reduced proliferation without causing significant cell death, implying a cytostatic rather than cytotoxic effect. To determine whether APT1 inhibition disrupts cell-cycle progression, we examined the distribution of cells across the cell-cycle phases. ML348 treatment significantly altered the distribution of cells across the cycle (two-way ANOVA interaction, *p* = 0.0001). The proportion of cells in G1 increased from 31.0% in control to 39.2% in ML348-treated cells, an 8.2 percentage point gain corresponding to a relative increase of ∼27% (*p* = 0.0028, Sidak’s test). This was accompanied by a reduction in the G2 population from 28.2% to 20.5% (–7.7 points; ∼27% relative decrease, *p* = 0.0046). The S phase decreased from 33.0% to 28.4% (– 4.6 points; ∼14% relative reduction), though this change did not reach statistical significance (*p* = 0.089) (Figure 7). These findings demonstrate that ML348 impairs cell growth partly by disrupting normal cell-cycle progression, leading to G1 accumulation and reduced entry into S/G2.

**FIGURE 7.**
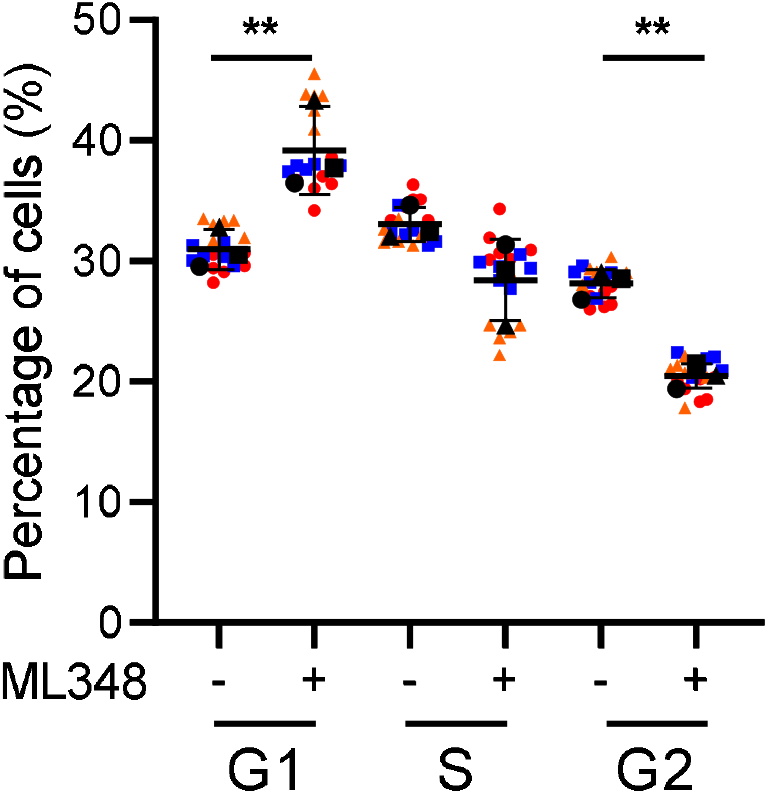
ML348 delays cell cycle progression. Cells were plated in 12-well plates and treated with 50 µM ML348 for 24 h. Following treatment, cells were fixed in 70% ethanol,stained with 0.1 µg/ml DAPI, and analyzed by flow cytometry. Cell cycle distribution was quantified using FlowJo. Statistical analysis was performed using two-way ANOVA with Sidak’s post-hoc test (**, *p* < 0.001). Data presented are mean ± SD (n = 3, with at least four technical replicates).

## Discussion

Our findings position *LYPLA1*/APT1 as a context-dependent vulnerability in breast cancer, driven primarily by enzymatic activity and subcellular localization rather than by expression level alone. Kaplan–Meier analysis showed that high *LYPLA1* expression correlates with poorer overall survival across nearly 5,000 patients (Figure 1A), consistent with emerging evidence that dysregulated protein *S*-acylation promotes tumor aggressiveness ^48–51^. Complementing this, large-scale CRISPR loss-of-function screens (DepMap) revealed variable reliance on LYPLA1 among breast cancer lines (Figure 1B), prompting us to investigate what defines APT1 dependency.

Comparative profiling of APT1 across molecular subtypes revealed that although luminal A MCF7 cells expressed the most protein (Figure 2), basal-like TNBC cells 468 displayed the highest specific enzymatic activity after normalization (Figure 3D). This decoupling of expression from function suggests that post-translational regulation or subcellular redistribution enhances APT1 activity in specific contexts ^23,30,52^. In particular, elevated mitochondrial APT1 activity in 468 cells (Figure 3B) raises the possibility that APT1 contributes to the metabolic reprogramming typical of aggressive TNBC subtypes, which depend heavily on mitochondrial metabolism ^53–55^. We identified several mitochondrial proteins with increased acylation following APT1 inhibition (Figure 5). A recent mouse model with liver-specific deletion of APT1 similarly revealed enrichment of mitochondrial proteins ^32^. Despite these molecular changes, APT1 deficiency did not alter mitochondrial mass or superoxide production in HepG2 cells, suggesting a subtle or context-dependent role for APT1 in mitochondrial regulation ^32^.

To explore how APT1 inhibition reshapes signaling, we profiled the *S-*acyl proteome following ML348 treatment. Global acyl-RAC analysis revealed no change in bulk *S-*acylation, implying that APT1 fine-tunes a limited set of substrates (Figure 4). This agrees with prior studies where the APT1/2 knockout caused no detectable global shift ^56^. The absence of detectable bulk changes in *S-*acylation may reflect several factors. First, functional redundancy among other cytosolic (e.g., ABHD17 isoforms ^57^, APT2 ^30^) and mitochondrial thioesterases (e.g., ABHD10 ^58^; ABHD11 ^59^) could mask the impact of APT1 loss. Second, pulse-chase experiments suggest that the kinetics of *S*-acylation turnover are slower than previously appreciated ^56^, potentially limiting the window in which deacylated substrates can be observed. Third, only a fraction of a given protein may be subject to *S-*acylation, making it difficult to detect changes amongst a large background of unmodified proteins. This may partly explain why many *S-*acylation studies focus on signaling proteins, where even a small shift in the proportion of modified protein can have significant downstream effects. In contrast, modifying a small fraction of a highly abundant metabolic enzyme may have subtler consequences, making such changes harder to detect or interpret. It may explain why APT1 effects on mitochondrial regulation are subtle or transient.

WST-8 assay showed a pronounced reduction in viable cell numbers after treatment, indicating that APT1 activity is required for normal proliferation (Figure 6A). However, apoptosis and necrosis remained minimal (Figure 6C, D), suggesting that this effect was cytostatic rather than cytotoxic, aligning with previous findings that APT1 knockout impairs tumorigenesis in TNBC (231) cells ^60^. To clarify how proliferation was suppressed, we performed cell-cycle analysis, which revealed a coordinated shift in phase distribution—an accumulation of cells in G1 accompanied by a complementary decrease in G2 (Figure 7). This pattern aligns with the proteomic evidence of increased acylation among replication and chromatin-associated proteins, suggesting that loss of APT1 activity disrupts the G1/S transition and slows cell-cycle progression. Together, these findings establish APT1 as a regulator of cell proliferation ^20^ through its control of key nuclear substrates involved in DNA replication and chromatin organization.

Our mass spectrometry analysis revealed increased S-acylation of several proteins typically associated with nuclear functions such as DNA replication and chromatin regulation (Figure 5). Among these, *MCM2* is a core component of the replicative helicase complex required for origin licensing and DNA unwinding ^42^, while *thymidine kinase 1 (TK1)* provides thymidine monophosphate for nucleotide synthesis ^43^. *DHX9* coordinates transcription and replication fork progression ^44^, and *RBBP4* and *RBBP7* participate in chromatin remodeling and transcriptional control ^45,46^. Curiously, our workflow typically removes the nuclei prior to analysis by Acyl-RAC. This raises the possibility that these proteins may undergo nucleo-cytoplasmic shuttling and be *S-*acylated in the cytosol. Similarly, as our cells are not synchronized for the cell cycle, a small percentage of the cells are likely in late prophase of mitosis after the nuclear envelope has been disassembled, raising the possibility that APT1 has a very specific function at this stage of the cell cycle. Ultimately, the enrichment of these proteins suggests that APT1 inhibition could indirectly influence nuclear pathways that sustain cell proliferation and division.

Together, our data supports a model in which APT1 sustains TNBC proliferation by maintaining the dynamic deacylation of nuclear and mitochondrial regulators that coordinate growth and G1/S transition. This has both mechanistic and therapeutic implications: targeting APT1 could selectively impair proliferation in aggressive breast cancers without broadly disrupting normal cellular function, since its substrates include replication and mitochondrial regulators essential for sustaining tumor proliferation but less critical for normal cell homeostasis.

## Abbreviations

Acyl-RAC: Acyl–Resin-Assisted Capture
APT1: Acyl-Protein Thioesterase-1
DAPI: 4′,6-Diamidino-2-Phenylindole
DMEM: Dulbecco’s Modified Eagle Medium
DMSO: Dimethyl Sulfoxide
DPP: Depalmitoylation Probe
EDTA: Ethylenediaminetetraacetic Acid
Ex/Em: Excitation/Emission
FBS: Fetal Bovine Serum
HEPES: 4-(2-Hydroxyethyl)-1-Piperazineethanesulfonic Acid
LC–MS/MS: Liquid Chromatography–Tandem Mass Spectrometry
LYPLA1: Lysophospholipase-1
MMTS: *S*-Methyl Methanethiosulfonate
PBS: Phosphate-Buffered Saline
PMSF: Phenylmethylsulfonyl Fluoride
SDS–PAGE: Sodium Dodecyl Sulfate–Polyacrylamide Gel Electrophoresis
TNBC: Triple-Negative Breast Cancer
zDHHC: Zinc Finger DHHC-Type Containing Protein

## Author Contributions

Conceptualization, M.S. and G.D.F.; Methodology, M.S., H.S.Z., and M.T.; Investigation, M.S., H.S.Z., and M.T.; Resources, S.P., R.S.K., and B.C.D.; Data curation, M.S.; Visualization, M.S.; Writing—original draft preparation, M.S.; Writing—review and editing, G.D.F., B.C.D., and M.S.; Supervision, G.D.F.; Project administration, G.D.F.; Funding acquisition, G.D.F. and B.C.D. All authors have read and agreed to the published version of the manuscript.

## Data Availability Statement

The mass spectrometry proteomics data have been deposited to the ProteomeXchange Consortium via the PRIDE ^61^ partner repository with the dataset identifier PXD069551. All other datasets analyzed during the current study are publicly available from established databases as described in the Materials and Methods section. Additional information is available from the corresponding author upon reasonable request.

